# *In silico* model of axonal pathfinding during spinal cord regeneration in zebrafish larvae

**DOI:** 10.64898/2026.04.17.719187

**Authors:** Oskar Neumann, Mathar Kravikass, Nora John, Rahul Gopalan Ramachandran, Paul Steinmann, Vasily Zaburdaev, Daniel Wehner, Silvia Budday

## Abstract

Functional spinal cord repair in zebrafish is governed by regeneration-favorable biochemical and mechanical cues within the lesion microenvironment. Alterations in extracellular matrix composition and stiffness are closely associated with axon regeneration. However, experimentally dissecting the interplay between mechanical signals and axonal regrowth *in vivo* remains technically challenging. Here, we present an agent-based modeling framework to simulate stiffness-mediated axonal growth trajectories across the lesion. We use this model to explore potential mechanisms underlying the characteristic growth patterns observed during zebrafish spinal cord regeneration. Computational predictions were qualitatively compared with confocal imaging data obtained from larval zebrafish. These phenomenological comparisons revealed a close agreement between simulated and experimentally observed axon growth, indicating that experimentally observed patterns could be governed by transient changes in the stiffness profile of the spinal cord and lesion microenvironment. Hence, our computational framework provides an *in silico* platform for investigating the role of mechanical cues in axon regeneration in the injured spinal cord.

## 1. Introduction

Following spinal cord injury (SCI), axon regeneration fails in humans and most other mammals but occurs robustly in zebrafish, enabling functional recovery [1, 2]. Emerging evidence indicates that differences in the mechanical properties of the local microenvironment contribute to the distinct regenerative capacity of mammalian and zebrafish central nervous system (CNS) axons. In zebrafish, spinal cord tissue transiently stiffens during the course of regeneration which is reflected by an increase in apparent modulus [3]. In rodents the tissue softens after injury [4, 5]. Moreover, experimental softening of zebrafish spinal cord lesions, through quantitative alterations of the extracellular matrix (ECM) composition, impairs regenerative outcomes [6]. However, mechanistically dissecting the interplay between mechanical cues and axon regeneration *in vivo* is technically challenging, and current evidence remains largely correlative.

Recent findings have begun to provide insights into how different mechanical properties of CNS tissue affect axonal growth [7]. A potential key factor lies in a phenomenon termed durotaxis [8, 9, 10, 11], which was first observed for fibroblasts [8]. Durotaxis is a mode of directed cell migration that arises from the integration of mechanosensing and mechanotransduction mechanisms, whereby cells detect spatial differences in ECM stiffness [12] and convert them into polarized cytoskeletal dynamics, adhesion turnover, and force transmission that bias movement along stiffness gradients [13]. Positive durotaxis is cell migration toward stiffer ECM, while negative durotaxis is migration toward softer ECM; both are guided by stiffness gradients. Different cell types (e.g., mesenchymal stem cells and collective cell systems) can undergo positive (from soft to stiff) or negative durotaxis (from stiff to soft) [14].

Although gel-based *in vitro* systems allow controlled manipulation of mechanical properties, they do not fully capture the complexity of the *in vivo* environment. On the contrary, separating mechanical cues from accompanying biochemical and structural changes *in vivo* remains challenging [15, 10]. Experimental evidence for how and when axons, e.g., in the regenerating zebrafish spinal cord, are guided by positive or negative durotaxis *in vivo*, remains scarce: NG108-15 neuron-like cells used in an *in vitro* model system were shown to be highly sensitive with regards to the steepness of the stiffness gradients, exhibiting positive durotaxis for shallow gradients and negative durotaxis for steeper gradients [16], while the *in vivo* embryonic *Xenopus* brain system for morphogenesis clearly shows a trend of axon bundles turning towards softer tissue when growing on a stiffness gradient [7]. Together, these findings suggest that axonal responses to mechanical heterogeneity are not governed by a single universal rule, but are likely shaped by the local combination of stiffness gradient magnitude, ECM composition and structure, and tissue architecture. *In silico* modeling is a valuable tool for disentangling these competing influences.

Here, we report on the development of a two-dimensional agent-based *in silico* model to assist the investigation of possible influences of stiffness-based axonal pathfinding modes in the context of spinal cord injury in zebrafish [3, 6, 17]. Based on experimental evidence showing that neurite growth is straighter in stiffer environments, while softer substrates promote more exploratory growth [7, 18, 19], the neuronal growth model is extended to incorporate stiffness gradient-guided axonal growth processes. This enables a computational investigation of neuronal growth patterns under a range of stiffness gradients and introduces a paradigm for probing transient, mechanical cues, which have been shown to dynamically guide collective cell migration by continuous remodeling of the surrounding tissue [20]. Our approach is flexible and versatile in terms of model configuration and general experimental parameters. The simulations are qualitatively validated against microscopic time-lapse live imaging of zebrafish larvae during regeneration, enabling a comparison between experimentally observed and simulated neuronal pathfinding behaviors.

## 2. Methods

### 2.1. Zebrafish husbandry and transgenic lines

Zebrafish lines were kept and reared at the Max-Planck-Zentrum für Physik und Medizin under a 14/10 h light-dark cycle at 28.5°C and according to FELASA recommendations, and the supervision of the veterinary authorities of the city Erlangen (permit: VII/390/DL003 of the Amt für Veterinärwesen und gesundheitlichen Verbraucherschutz Stadt Erlangen). All experimental procedures were done on larval zebrafish aged up to 120 hours post-fertilization (hpf), derived from voluntarily mating adult zebrafish. Procedures on zebrafish larvae up to 120 hpf are not regulated as animal experiments by the European Commission Directive 2010/63/EU. For all experiments, we used Tg(*elavl3* :rasmKate2)^mps1^ and Tg(*elavl3* :GFP-F)^mps10^ transgenic zebrafish lines [6, 21].

### 2.2. Zebrafish spinal cord lesions

A detailed protocol for inducing spinal cord lesions in zebrafish larvae has been previously described [22]. Briefly, zebrafish larvae (3 dpf) were anesthetized in E3 medium containing 0.02% MS-222 (PharmaQ, Cat. # Tricaine PharmaQ). A 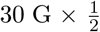 in hypodermic needle was used to transect the spinal cord by either incision or perforation at the level of the urogenital pore. After surgery, larvae were returned to E3 medium for recovery and maintained at 28.5 °C.

### 2.3. Live imaging of larval zebrafish

For live confocal imaging, zebrafish larvae were anesthetized in E3 medium containing 0.02% MS-222 and mounted in the appropriate orientation in 1% low-melting-point agarose (Ultra-Pure™ Low Melting Point, Invitrogen, Cat.#16520) between two microscope cover glasses. During imaging, larvae were covered with E3 medium containing 0.01% MS-222 to prevent the preparations from drying out. Imaging was performed using a Plan-Apochromat 20×/0.8 objective on a Zeiss LSM 980 confocal microscope.

### 2.4. Image analysis

We increased the intensity of the acquired microscopic images by 20% to improve the visibility of neuronal cells using ImageJ (https://imagej.net/ij/; version 1.54p).

### 2.5. Agent-based modeling

The agent-based model used in this study builds on the *in silico* neuritogenesis framework originally developed by Kravikass et al. [19] and first applied by Furlanetto et al. [23] to describe neurite outgrowth dynamics in the context of neurodevelopmental disorders. In this coarse-grained, two-dimensional model, neurites are represented as chains of circular beads connected by harmonic springs, protruding from an immobile cell body. The leading bead of the chain represents the growth cone and drives extension. The growth cone selects a random ECM particle within a defined cone angle and range ahead of the neurite tip and forms a temporary adhesive link modeled as an elastic spring. This link generates a traction force between the growth cone and the ECM particle. The link persists for a characteristic time before breaking, after which a new ECM particle is selected and the process repeats. When the distance between the growth cone and the preceding bead reaches a critical length, a new bead is inserted into the chain, elongating the neurite.

The ECM is modeled as a collection of circular particles. In the configuration used here, the ECM particles are connected by elastic springs via Delaunay triangulation. All particles interact through excluded volume forces that prevent overlap. The equations of motion are solved in the overdamped limit, where inertial terms are neglected and the displacement of each particle at every time step is determined by the balance of spring, traction, lattice, and excluded volume forces, scaled by the respective particle mobility.

Key parameters governing growth behavior include the cone angle, which controls directional persistence, the link range, the link persistence time, and the link spring constant. The model is formulated in dimensionless form and can be rescaled to match experimental observations by calibrating spatial and temporal reference scales against known physical quantities. The corresponding simulation parameters used here can be found in Tab. S1 and Tab. S2.

#### 2.5.1. Implementation of mechanosensing

To incorporate the ability of the growth cone to “sense” the mechanical properties of its local environment and correspondingly modulate growth-specific parameters, we allow two parameters of the original model ([19, 23]) to vary in response to the local ECM stiffness. With this, the ability of axons growing straighter on stiffer substrates and exhibiting more exploratory, less persistent growth on softer substrates is probed in a much broader range than what is possible due to passive mechanical interactions [19]. In addition, we allow growth speed to be coupled to environmental stiffness, such that growth is slower in softer and faster in stiffer environments. Mathematically, we describe both mechanisms based on a mechanosensing weight for each neurite’s leading bead which is computed every time a new link between a neurite leading bead particle and an ECM particle is created in the agent-based simulation. This weight quantifies the local environment stiffness of an individual neurite leading bead in the following procedure:

Let the leading bead particle be at position **x**_t_ ∈ ℝ^2^. All ECM particles *p* within a mechanosensing range *R*_ms_ form the set of neighbouring ECM particles

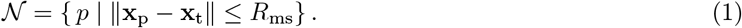

For each *p* ∈ 𝒩, let 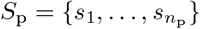 be the set of springs associated with *p* (see Fig. 1a). Each spring *s* ∈ *S*_p_ is mechanically characterized by a spring constant *k*_s_. Letting *n*_p_ := #𝒩 denote the size of the set 𝒩, the mean spring constant of the ECM particles connected to particle *p* (see Fig. 1b; exemplary springs for one neighbouring ECM particle shown in red) is given by

**Figure 1:**
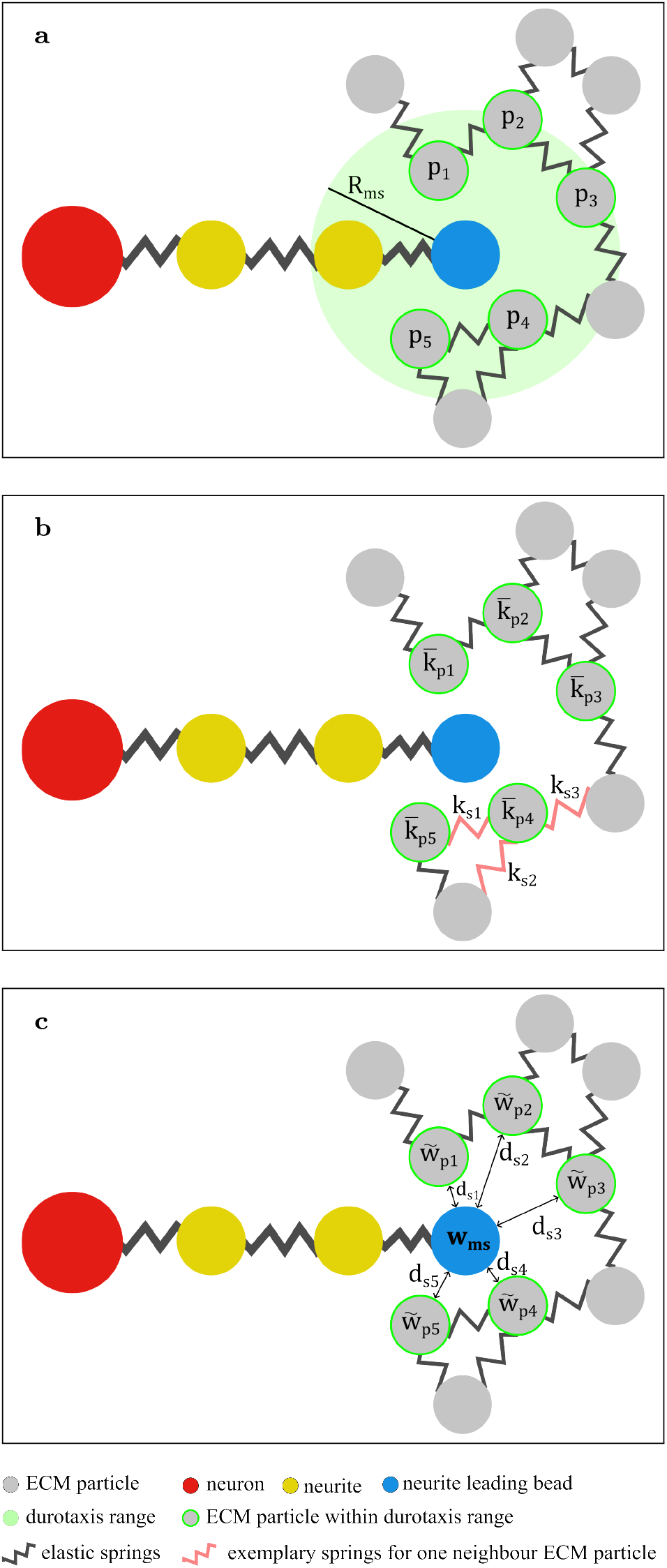
**(a)** Schematic illustration of the mechanosensing range around a neurite leading bead, including all neighboring extracellular matrix (ECM) particles within this range. **(b)** Schematic illustration of the averaging procedure used to compute the effective spring coefficient from the surrounding ECM particles. Red springs indicate the connections utilized in Eq. 2 when computing the mean spring constant of 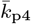. **(c)** Schematic illustration of the computation of mechanosensing weighting. Variables defined in Eqs. 1–6

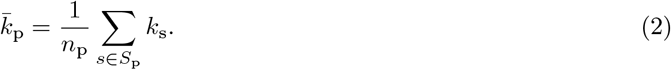

The raw weight per particle within *R*_ms_ is scaled by the inverse of the distance to the neurite leading bead, based on the assumption that the further away a particle, the smaller its contribution to the stiffness sensed by the leading bead. The raw and normalized weights are thus

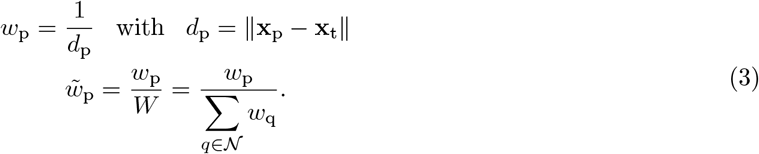

Finally, the averaged normalized environment stiffness sensed at the tip (see Fig. 1c) is calculated relative to the maximum spring constant, *k*_max_, in the entire spring lattice of the simulation domain:

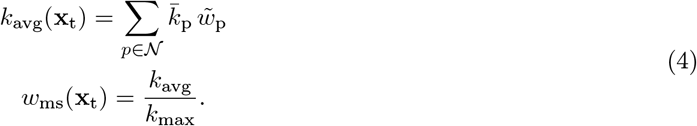

The growth rate is then coupled to the environment stiffness sensed by an advancing neurite leading bead via the link range which defines the radial extent of the search cone in which an ECM link will be selected for a new neurite-ECM link (see Fig. 2a). The higher the distance between ECM particle and neurite leading bead particle, the higher the resulting spring force, thus the higher the rate at which leading bead neurites move per time step. For this, the initial link range *R* is scaled using a logistic function with midpoint at 0.5 and slope of *β*_link_ thereby effectively increasing local neurite growth speed in stiffer environments in a controlled fashion.

**Figure 2:**
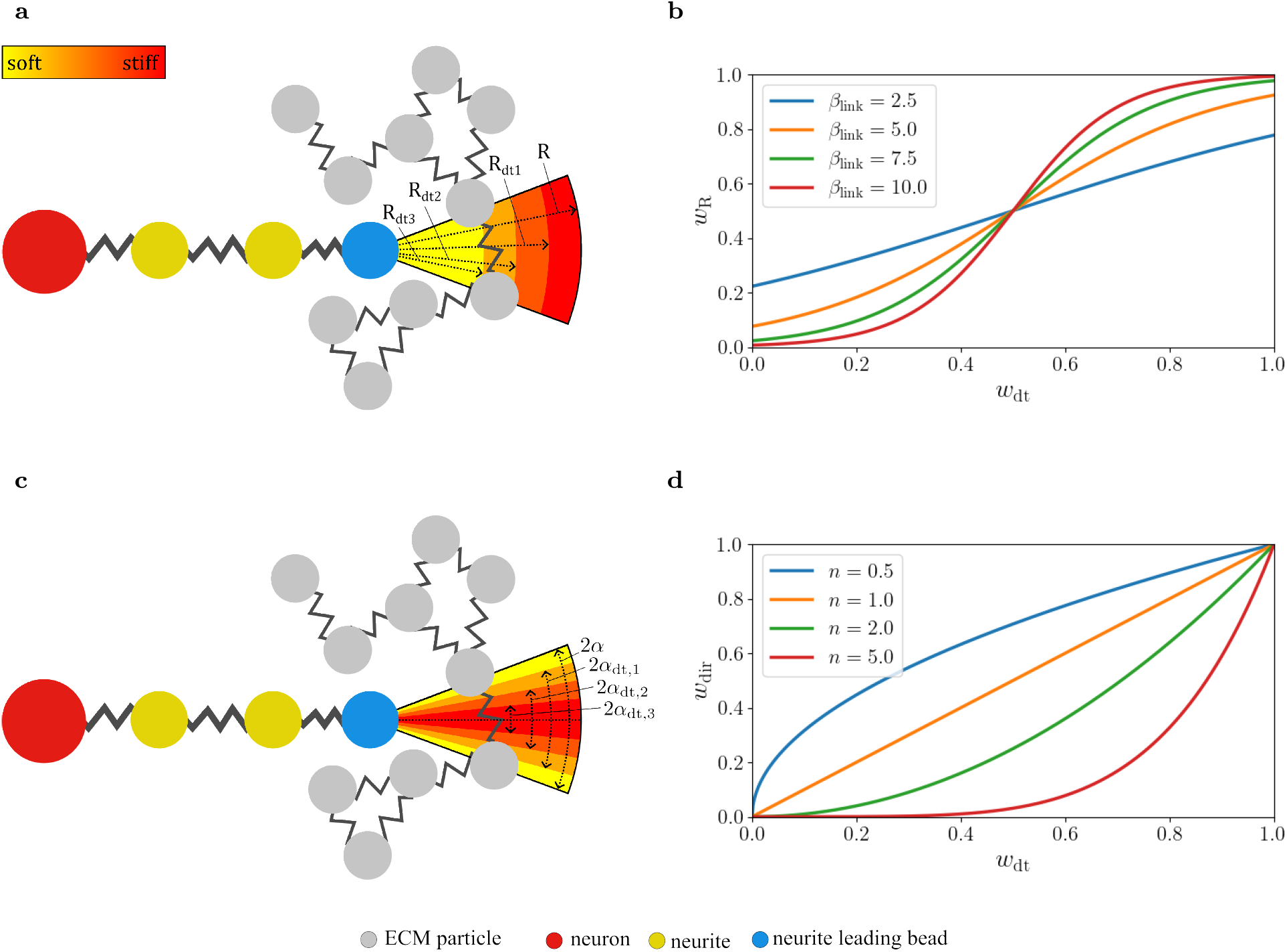
Illustrations of mechanisms of mechanosensing within our modeling approach with neurons, neurites forming an axonal chain, neurite leading beads and ECM particles as red, yellow, blue and grey circles. Connections between particles are visulized by springs. **(a)** Schematic illustration of the stiffness-dependent link range around the neurite leading bead. *R* refers to the initial link range and *R*_dt1_, *R*_dt2_ and *R*_dt3_ to weighted link ranges for decreasing environment stiffness values. **(b)** Corresponding mechanosensing weighting curves for different values of the parameter *β*_link_ based on Eq. 5. *α* refers to the cone angle and *α*_dt1_, *α*_dt2_ and *α*_dt3_ to weighted cone angles for increasing environment stiffness values. **(c)** Schematic illustration of the stiffness-dependent cone angle defining the directional sensing region *α* refers to the initial cone angle and *α*_dt1_, *α*_dt2_ and *α*_dt3_ to weighted cone angles for decreasing environment stiffness values. **(d)** Corresponding mechanosensing weighting curves for different values of the parameter *n* based on Eq. 6.

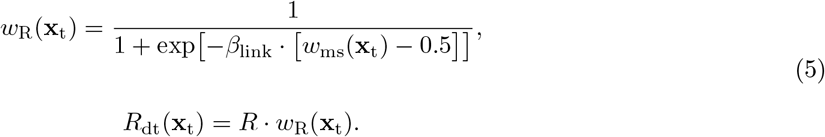

Similarly, the initial cone angle *α*, which defines the angular range over which a neurite leading bead searches for a new ECM particle to connect to, is scaled by a power law with exponent *n*. This scaling amplifies the already present “passive” persistence mechanism in the original model framework [19] and introduces a controlled stiffness-dependent directionality in the search for a new ECM-link partner (see Fig. 2c). Consequently, a higher average environmental stiffness narrows the angular range of the growth cone, promoting straighter axon growth, whereas lower stiffness leads to broader, exploratory growth.

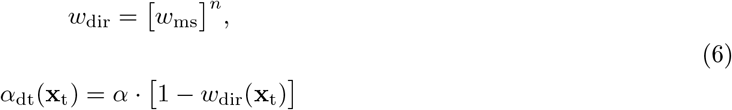

Fig. 2b and c highlight the attenuation of the link range and cone angle by different values for the corresponding mechanosensing parameters *β*_link_ and *n*.

As a summary, the parameters governing axon growth patterns in the presented model include the minimum and maximum lattice stiffness, *k*_min_ and *k*_max_, which define the range of spring coefficients from which the lattice is constructed; the mechanosensing range *R*_ms_, which determines the circular area in which neighboring ECM particles and their elastic springs are mechanically sensed by a growth cone; the stiffness-based growth speed parameter *β*_link_, which reflects the sensitivity of the growth speed to the local environmental stiffness; and the stiffness-based directionality parameter *n*, which captures the relationship between environmental stiffness and axon growth directionality.

#### 2.5.2. Realization of stiffness gradients

As described above, the environmental stiffness sensed by a neurite leading bead is defined as the normalized, locally averaged spring coefficient of neighboring ECM particles. Accordingly, in the following analyses, stiffness gradients within the ECM are interpreted and visualized by scatter plots of ECM particles, where color encodes the locally averaged spring coefficient of individual particles. Test cases were simulated with an initial growth cone angle of *α* = 30°. The progressive wound edge cases were simulated with an initial growth cone angle of *α* = 30° and *α* = 75°.

#### 2.5.3. Test cases

The stiffness of the extracellular environment in the spinal cord and at the lesion site is complex and spatially heterogeneous, owing to variations in tissue composition and differences between white and gray matter. It is therefore crucial to assess how the presented model responds to both abrupt and gradual changes in stiffness.

For the construction of stiffness gradients, the spring coefficients of the ECM lattice were assigned values within a predefined interval between *k*_min_ and *k*_max_, which were kept fixed and consistent across all simulations in this study (see Tab. S1). The following test scenarios were selected to evaluate the simulation

A banded stiffness gradient (Fig. 3a) was implemented along the vertical axis of the simulation domain by dividing the domain height into horizontal bands of fixed thickness. Each band was assigned a uniform stiffness value, increasing stepwise from *k*_min_ and *k*_max_ from top to bottom boundary. In this way, the stiffness varied monotonically along the vertical direction while remaining constant within each band, yielding a piecewise-constant approximation of a linear stiffness gradient.

**Figure 3:**
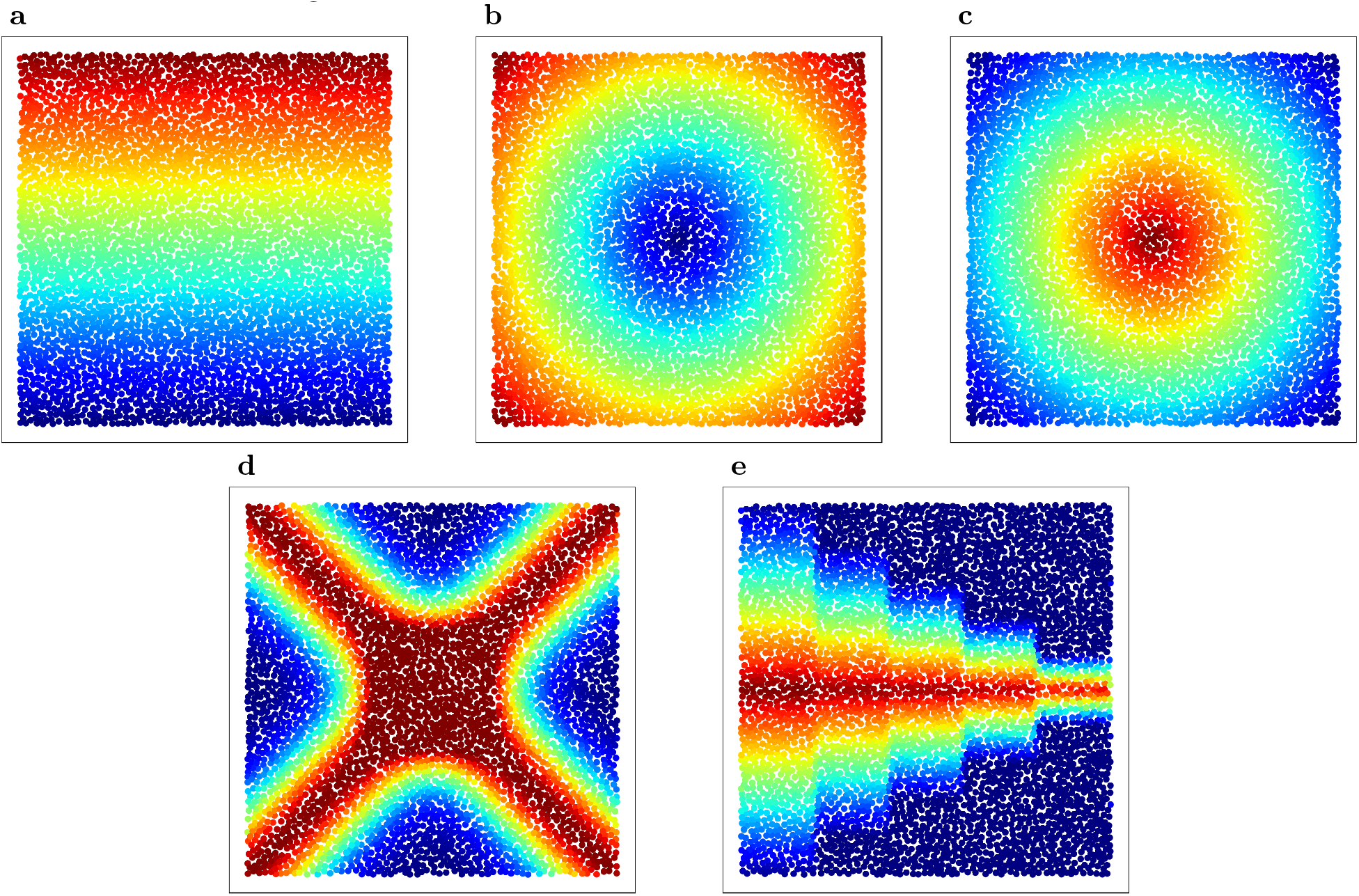
ECM lattice stiffness distributions for (a) a horizontal band pattern, (b) a radially increasing pattern with a soft centre, (c) a radially decreasing pattern with a stiff centre, (d) a cross-shaped pattern with stiff diagonal bands, and (e) a cascading band pattern. Colours indicate local stiffness, with blue denoting softer regions and red denoting stiffer regions.

For the radial stiffness configuration, the local stiffness was defined as a monotonically increasing (Fig. 3b) or decreasing (Fig. 3c) function of the distance from the center of the simulation domain. Regions closer to the center were assigned higher stiffness values, with stiffness gradually decreasing toward the periphery and remaining bounded between *k*_min_ and *k*_max_ in Fig. 3b; the opposite configuration was used for Fig. 3c.

A cross-shaped stiffness pattern (Fig. 3d) was defined by superimposing two diagonal stiff bands that intersect at the center of the simulation domain. The bands follow the two lines passing through the domain center with slopes ±*m*, where *m* = *L*_y_*/L*_x_ for a domain of size (*L*_x_, *L*_y_). For a spatial position (*x, y*), we computed the perpendicular distances *d*_1_ and *d*_2_ to these two lines and mapped each distance to a Gaussian band profile 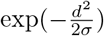 with a width parameter *σ* = 0.1· *L*_y_. The two contributions were summed and capped to unity to obtain a normalized value, which was then used to interpolate linearly between the prescribed stiffness bounds *k*_min_ and *k*_max_. Consequently, stiffness attains values close to *k*_max_ along the two diagonals and smoothly decays toward *k*_min_ away from the bands.

A cascaded band pattern (Fig. 3e) was implemented by combining a global vertical band profile with stripe-wise modifications along the horizontal axis. The domain was first normalized in the horizontal direction as *x*_frac_ = *x/L*_x_. For *x*_frac_ ≥ 0.2, the domain was subdivided into vertical stripes of width 0.2 · *L*_x_ and indexed by

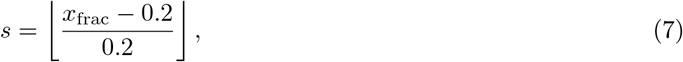

with an associated cut fraction

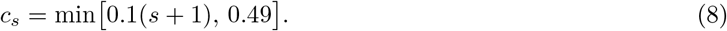

Within each stripe, only a central corridor in the *y*-axis was retained,

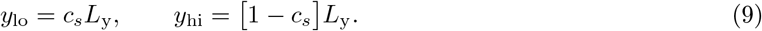

and points outside this corridor were assigned *k*_min_. Inside the corridor, stiffness increased stepwise from the corridor edges toward the corridor centerline using horizontal bands of fixed thickness. Specifically, defining the distance to the nearest corridor edge

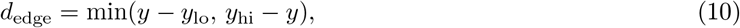

stiffness was mapped linearly from *k*_min_ at *d*_edge_ = 0 to *k*_max_ at the corridor midline using banded discretization. Outside the stripe region (*x*_frac_ *<* 0.2), the following pattern was applied: stiffness was maximal near the center midline (*y* = *L*_y_*/*2) and decreased stepwise toward the top and bottom boundaries.

#### 2.5.4. Progressing wound edge

A transient wound-edge stiffness pattern (Fig. 4a–c) was generated by superimposing two elliptical contributions located at the left and right boundaries of the domain. Two ellipse centers were defined at

**Figure 4:**
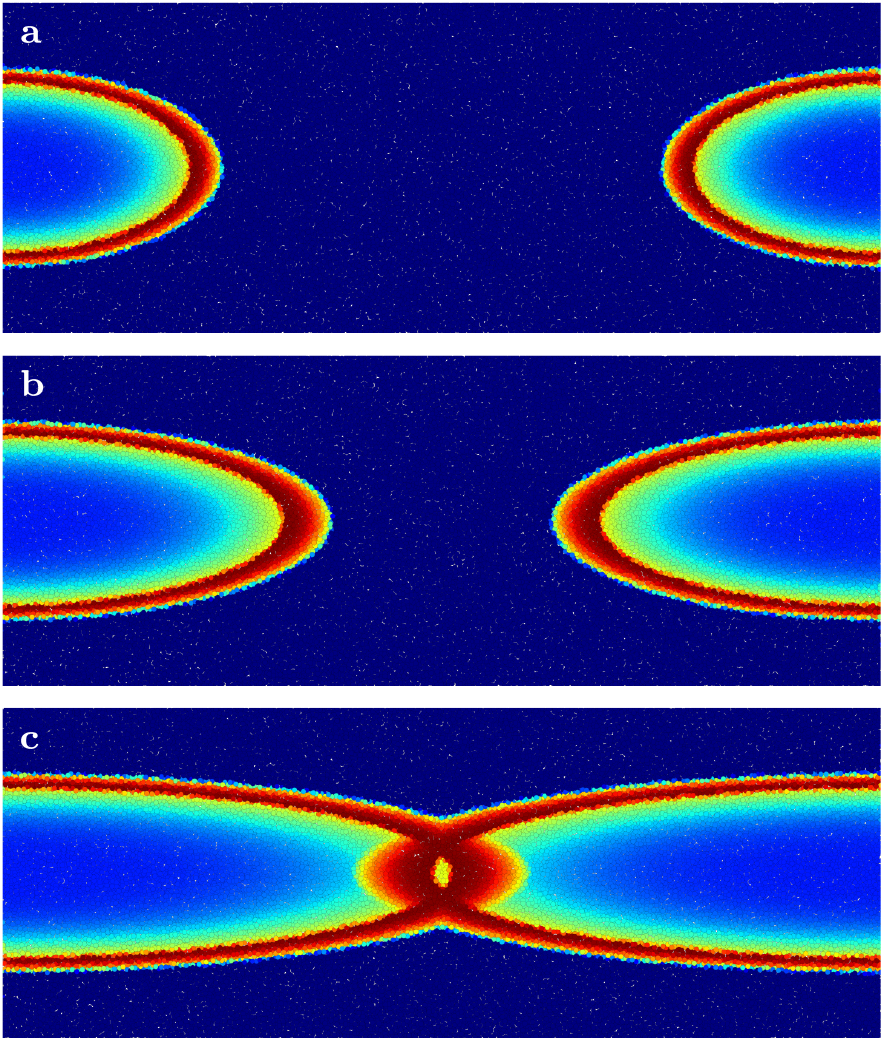
**(a)** Stiffness gradient representing the wound edge during the initial simulation phase (*t*_sim_ *<* 0.25 *t*_end_). **(b)** Stiffness gradient representing the wound edge during the intermediate simulation phase (0.4 *t*_end_ ≤ *t*_sim_ *<* 0.6 *t*_end_). **(c)** Stiffness gradient representing the wound edge during the late simulation phase (*t*_sim_ ≥ 0.6 *t*_end_). Colors indicate local stiffness, with blue denoting softer regions and red denoting stiffer regions.

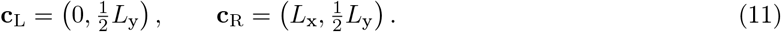

with semi-axes *R*_*x*_ based on three simulation steps through the course of computation:

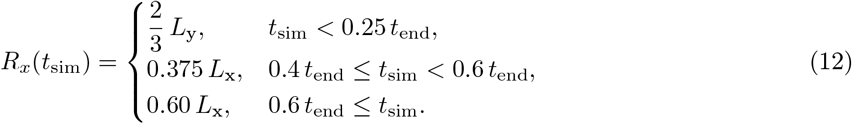

The vertical semi-axis was constantly fixed at *R*_*y*_ = 0.3 *L*_y_. For a position **p** = (*x, y*), we computed the normalized elliptical radius

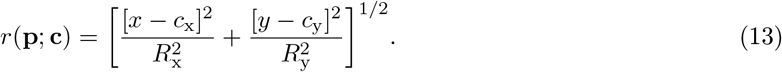

for **c** ∈ {**c**_L_, **c**_R_}, such that *r* = 1 corresponds to the ellipse boundary. The local stiffness contribution of one ellipse was then defined as a piecewise function of *r*: outside the ellipse (*r* ≥ 1) stiffness was set to *k*_min_; inside the ellipse (*r <* 1) a narrow, stiff rim was formed near the boundary using a transition radius *r*_*b*_ = 0.85 and power-law shaping with exponents *p*_in_ = 4 and *p*_out_ = 3. With 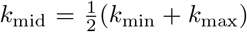, this was implemented as

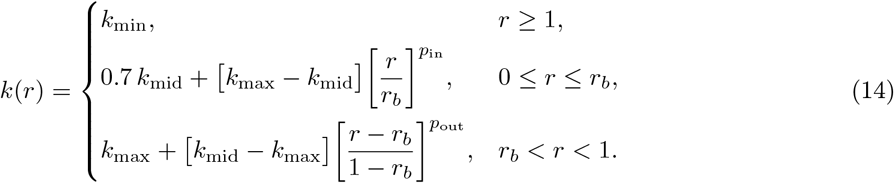

Finally, the wound-edge stiffness field was defined as the maximum of the left and right contributions,

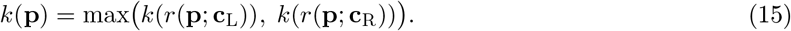

The parameters introduced above (dimensions of the wound edge, the time points at which it evolves, the transition radius, and the power-law exponents) were selected to produce the desired qualitative behavior in the simulations. Using three distinct values for the horizontal semi-axis, the progressively evolving stiff wound edge was discretized into three successive stiffness gradients, as illustrated in Fig. 4a–c.

For simulations involving progressively evolving wound-edge stiffness gradients, in each simulation, the growth of 26 individual axons was initiated at the left boundary of the domain, representing the rostral side.

### 2.6. Statistical analysis of neurite growth

Following [23], for each trajectory *j*, representing a grown axon, the path length traveled up to time *t* = *N* · *dt* (after *N* time steps) was computed as

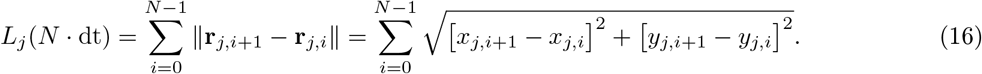

and the corresponding end-to-end distance as

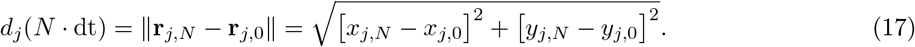

Tortuosity was defined as the ratio

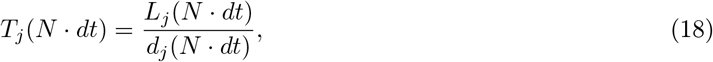

where *T* = 1 corresponds to a straight trajectory, and ensemble averages were obtained by averaging over all trajectories available at time *N* · *dt*. The mean squared displacement (MSD) was computed according to

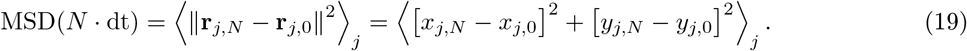

with ⟨·⟩_*j*_ denoting an average over multiple trajectories within one simulation and across simulations with the same parameter sets (*β*_link_, *n, α*).

Trajectories or axons were classified as crossing if they terminated at positions with *y <* 0.5 · *L*_y_ after originating from *y >* 0.5 · *L*_y_, or vice versa, and regeneration was considered complete when axons reached positions with *x >* 0.9 · *L*_x_.

For each parameter set of (*β*_link_, *n, α*) five simulations were performed and mean and standard deviations were computed.

## 3. Results

### 3.1. Morphological observations of axon regeneration in the larval zebrafish spinal cord

Fig. 5 (dorsal view) and in Fig. 6 (lateral view) present time-lapse confocal images of the emerging patterns of axonal regrowth after spinal cord transection in zebrafish larvae [24]. Fish 1 in Fig. 5 shows that individual axons either grow in seemingly random direction, in some cases even perpendicular to the rostro–caudal axis without any chance of crossing the lesion core (Fish 1, third and fourth rows from the top; Fish 5, bottom row), or, to a greater extent, at smaller angles to the rostro–caudal axis crossing the gap between the two healthy stumps. The dorsal series of Fish 1 suggests a characteristic reorientation behavior: axons appear to undergo discrete changes in growth direction within the lesion core, producing crossing, X-like trajectories and culminating in a horizontally aligned, hourglass-like axonal architecture at 48 hours post lesion (hpl). A similar morphology is evident at 48 hpl in both Fish 1 and Fish 2 in Fig. 5, indicating that this crossing pattern may represent a recurrent endpoint of regeneration in the dorsal projection.

**Figure 5:**
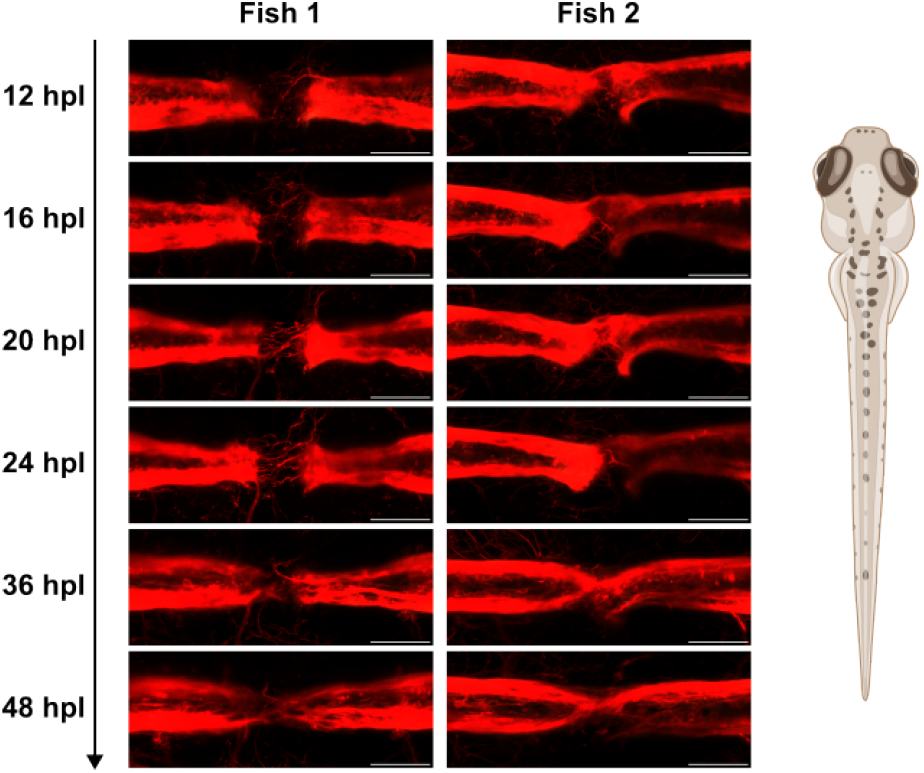
Dorsal-view time-lapse confocal images of axonal regrowth during spinal cord regeneration of zebrafish larvae. Maximum-intensity projections (medio–lateral) are shown at 12, 16, 20, 24, 36, and 48 hpl (from top to bottom). Trajectories represent 2D projections of 3D paths. Scale bars: 50 *µ*m. The dorsal-view zebrafish sketch was created using BioRender.

**Figure 6:**
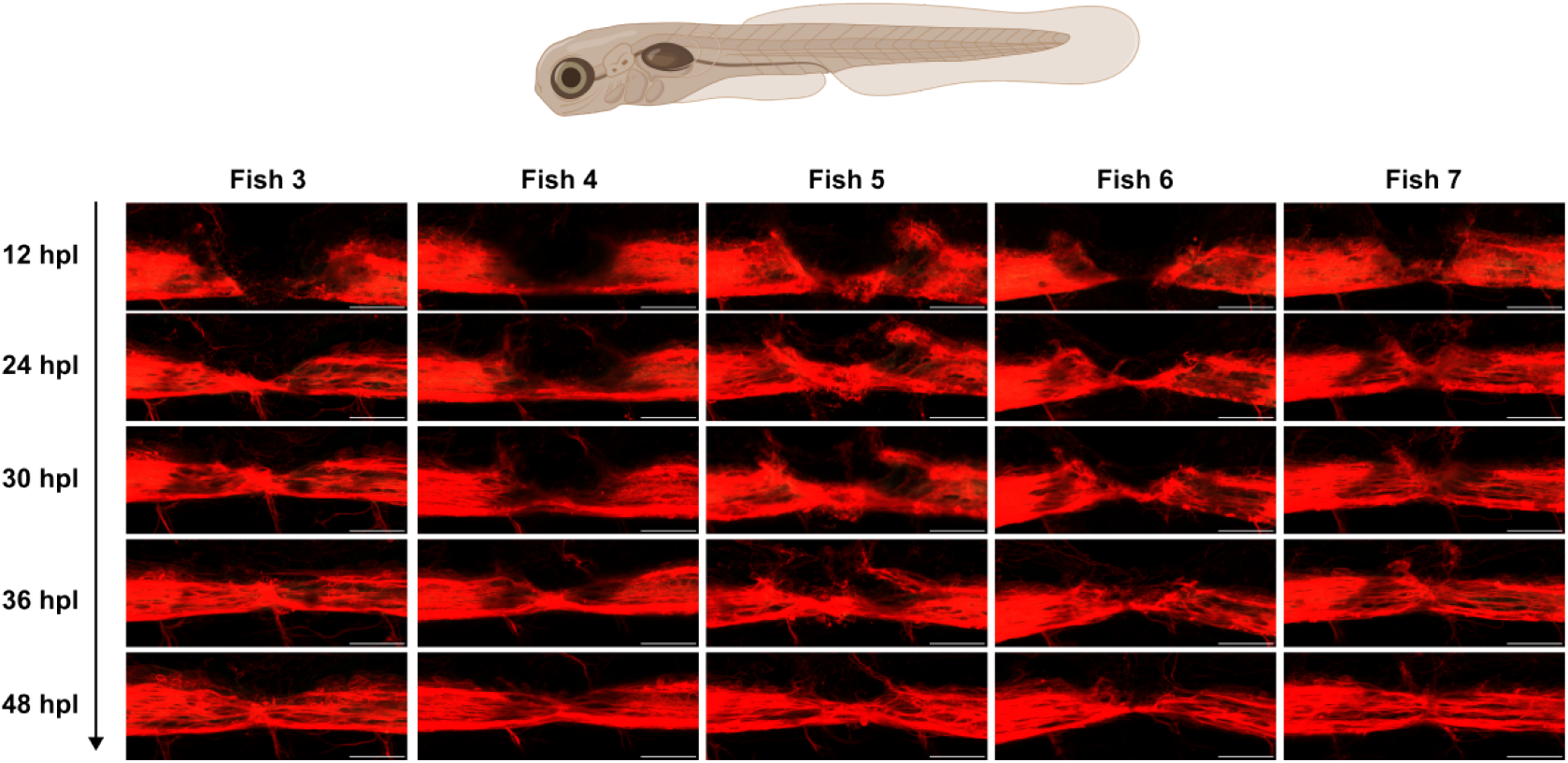
Lateral-view time-lapse confocal images of axonal regrowth during spinal cord regeneration of zebrafish larvae. Maximum-intensity projections (dorso–ventral) are shown at 12, 24, 30, 36, and 48 hpl (from top to bottom). Trajectories represent 2D projections of 3D paths. Scale bars: 50 *µ*m. The lateral-view zebrafish sketch was created using BioRender.

The lateral-view time-lapse images in Fig. 6 reveal a comparable X-shaped neuronal growth pattern at 48 hpl in Fish 3, Fish 4, Fish 7 and to a lesser extent in Fish 5 and Fish 6. Across all specimens, the apparent regrowth dynamics show a weaker progression from the lateral margins of the lesion towards the dorsal side. In parallel, the junction point of the re-established axonal bridge, which reconnect the severed spinal cord stumps, appears to shift dorsally over time and, in Fish 3 and Fish 4, becomes approximately centered along the dorso–ventral axis by 48 hpl. Fish 5 exhibits a highly disorganized axon arrangement at 12 hpl that progressively consolidates during regeneration, resulting in an hourglass-like architecture at 48 hpl that resembles the patterns observed in the other biological replicates. In contrast, Fish 6 shows an initially tilted orientation of both rostral and caudal stumps towards the dorsal side at 12 hpl, which results in a less stereotypical endpoint morphology at 48 hpl; nevertheless, a constricted bridging region around the lesion center remains evident. Fish 7 largely recapitulates the regrowth sequence observed in Fish 3 and Fish 4. Consistent with the dorsal-view series (Fig. 5), the lateral-view time-lapse images in Fig. 6 suggest discrete reorientation events of individual axons as they approach and traverse the lesion core. In Fish 3, 4 and 7, the final neuronal architecture at 48 hpl is compatible with a dorso–ventral exchange of trajectories across the lesion, i.e., axons that appear dorsal on the rostral side, project to more ventral positions on the caudal side, and vice versa.

### 3.2. Agent-based in silico modeling of axonal path finding

In the following, we use the agent-based model (ABM) introduced in Section 2.5 to explore possible relationships underlying the experimental observations presented in Section 3.1 by isolating the direct effects of different mechanisms of mechanosensing on the simulated axonal growth patterns.

#### 3.2.1. Influence of mechanosensing on axonal growth patterns

To qualitatively explore the effect of different mechanisms of mechanosensing, we use our ABM approach and analyze the observed growth characteristics of axons during successful regeneration in the zebrafish larvae *in silico*. Fig. 7 shows neuronal growth of 25 individual neurons for five different stiffness gradients and different stiffness-dependent growth parameters. Fig. 7, second column, shows the results for neurons growing just by the rule set of the original ABM [19], which recapitulates the effect of an increased persistence in stiffer and more dense ECM. However, for a broad range of parameters tested, we do not observe any pronounced effects of introduced stiffness gradients on the growth (see Fig. S1) patterns of neurites (see Fig. 7, column 2 for a representative image). Fig. 7, third column, shows the influence of coupling the link range to the environment stiffness, thus incorporating faster growth on stiffer substrates. For a better visibility of the desired effects of this coupling, the simulated trajectories are additionally forced to evolve straight. Fig. 7, fourth column, shows the influence of coupling the cone angle to the environment stiffness, allowing for growth on soft substrates to evolve in more tortuous paths, whereas growth on stiff substrates evolves more straight. Fig. 7, last column shows the combined effects of both stiffness-dependent growth speed and directionality.

**Figure 7:**
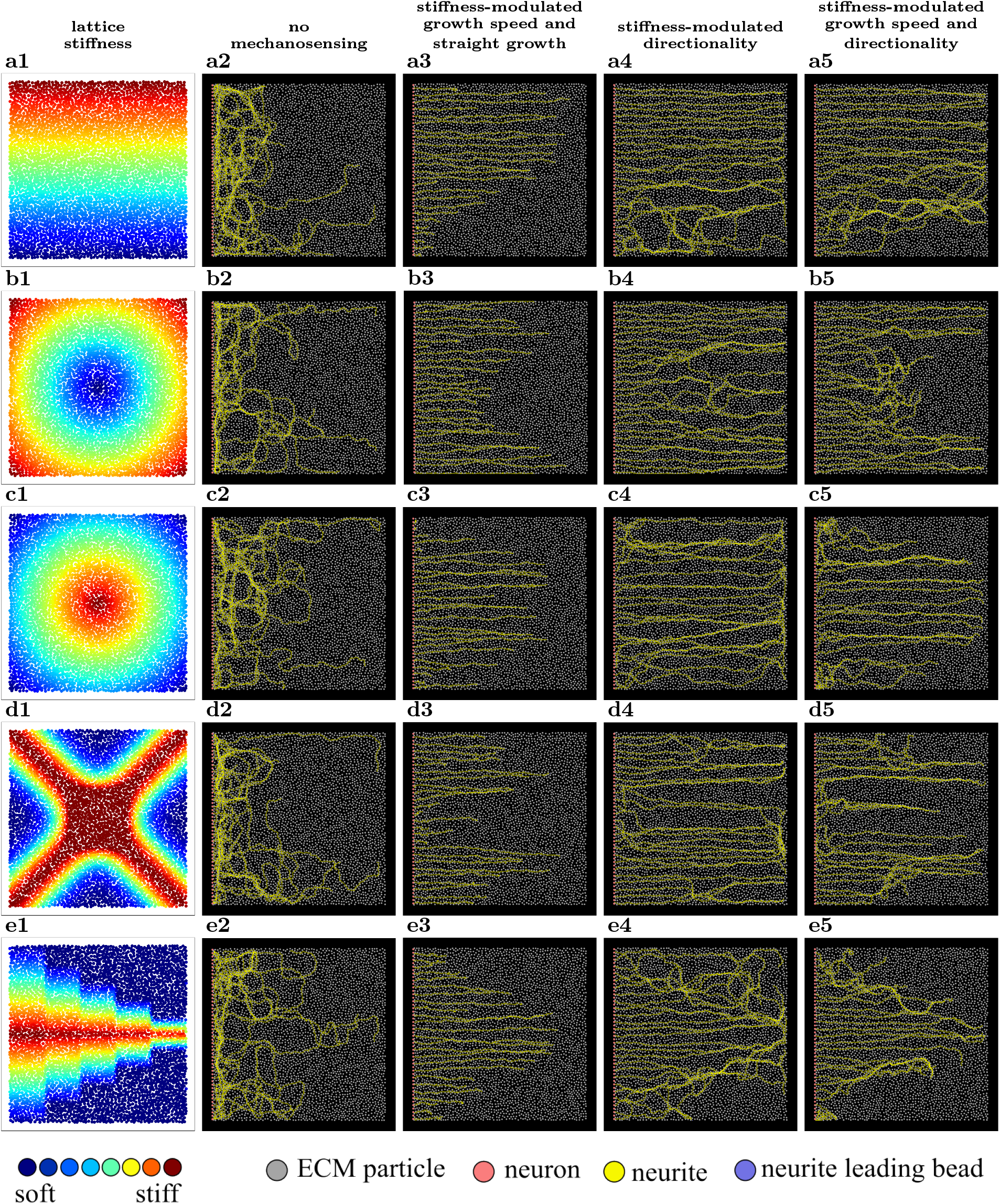
Overview of simulated test cases used to qualitatively assess mechanosensing-enhanced axonal growth in the agent-based model. Rows (a–e) correspond to different stiffness gradient configurations, including banded, radial (soft and stiff center), cross-shaped, and cascaded band patterns. The first column (a1–e1) shows the underlying ECM lattice stiffness distributions. The second column (a2–e2) displays axonal growth in the absence of mechanosensing, governed solely by the baseline particle-based growth rules. The third column (a3–e3) illustrates the effect of stiffness-modulated growth speed by coupling the link range to local stiffness, resulting in increased growth velocities on stiffer substrates; for clarity, growth is constrained to straight trajectories. The fourth column (a4–e4) shows the effect of stiffness-modulated directionality by coupling the growth cone angle to local stiffness, leading to more tortuous growth on soft substrates and straighter growth on stiff substrates. The fifth column (a5–e5) presents the combined effect of stiffness-modulated growth speed and directionality. The legend below the last row indicates particle types and the color coding of lattice stiffness, ranging from soft (blue) to stiff (red).

The simulated neuronal trajectories for the stiffness gradient with a gradually decreasing ECM stiffness from top to bottom (Fig. 7a1) show highly unorganized growth for the case without mechanosensing (Fig. 7a2). When the stiffness-modulated growth speed is included (Fig. 7a3), we observe the expected behavior, with substantial axonal growth of axons almost in the upper part of the domain, but little to no growth in the lower part of the problem domain. In results for stiffness-modulated directionality (Fig. 7a4), the upper half of the problem domain shows straight growth, while axonal growth shows more and more exploratory growth towards the softer bottom edge of the domain. When both mechanosensing effects are combined (Fig. 7a5), no axon is able to reach the right side of the domain on the bottom right area and we observe a strong tendency of axons growing towards the stiffer region in the lower part of the domain.

The trajectories for the radial ECM stiffness pattern with a soft core and stiff outer rim (Fig. 7b1) without mechanosensing show highly disorganized growth patterns, with individual axons growing at the top and bottom areas, avoiding to penetrate the center of the problem domain (Fig. 7b2). Including stiffness-modulated growth speed (Fig. 7b3) leads to faster axonal growth at the top and bottom, where the stiffness on average is highest, and slower growth in the softer center. Stiffness-modulated directionality during growth (Fig. 7b4) increases the degree of disorganization for growth through the softer center, while axons near the top and bottom of the problem domain grow straight. The combined effects of both stiffness-modulated parameters (Fig. 7b5) show no growth through the soft center but full outgrowth to the right side of the domain near the top and bottom of the domain.

The inverted radial ECM stiffness pattern with a stiff core and soft outer rim (Fig. 7c1) interestingly produces similar growth patterns to the test case before, when mechanosensing is disabled (Fig. 7c2). In line with previous results, the coupling of growth speed with ECM stiffness yields enhances growth in stiff areas around the center of the domain and inhibits growth near the top and bottom of the domain (Fig. 7c3). In contrast to the previous test case, for the stiffness-modulated directionality, we now observe straight growth near the middle of the problem domain and more tortuous growth near the top and bottom edges (Fig. 7c4). The combined effects of mechanosensing show a V-shaped pattern with almost full outgrowth to the right side of the domain in the middle part and gradually decreasing outgrowth from the center towards the top and bottom edges of the domain (Fig. 7c5).

The stiff X-shaped ECM stiffness pattern (Fig. 7d1) again shows a similar growth pattern to the two previous test cases without the influence of mechanosensing (Fig. 7d2). When stiffness-modulated growth speed is included, the fastest growth appears in two bands which seem to recreate the path of highest average stiffness across the stiffness gradient pattern at around 25 and 75 % of the total height of the problem domain (Fig. 7d3). When stiffness-modulated directionality is enabled, axon growth patterns are straight in the same regions of highest average stiffness and the most undirected in the horizontal center band of the domain (Fig. 7d4). The combined effects of both stiffness-modulated parameters superimpose both of the previously described growth patterns, with no axon being able to pass the center of the problem domain, leaving the small bands at 25 and 75 % of the total height as pathways for successfully crossing the whole problem domain during axonal growth (Fig. 7d5).

Lastly, the cascading ECM stiffness gradient with a growing softer domain at the top and bottom of the problem domain along five vertical bands (Fig. 7e1) results in highly disorganized growth near the top and bottom edges and more straight growth along the stiff centered band without mechanosensing (Fig. 7e2). Enabling stiffness-modulated growth speed shows the expected result of faster growth in the center and gradually decreasing outgrowth towards the bottom and top edges of the problem domain (Fig. 7e3). Stiffness-modulated directionality yields a similar result, although some more randomly growing axons at the top and bottom edges manage to cross the whole problem domain (Fig. 7e4). Finally, under the combined effects of stiffness-modulated growth speed and directionality, the growing axons form a vertical V-shape configuration, where all axons in the horizontal center band reach the right side of the domain, while horizontal axonal growth distances gradually decrease towards the top and bottom edges (Fig. 7e5).

These results demonstrate that the model incorporating both mechanosensing mechanisms yields the most pronounced response to stiffness patterns. We therefore focus on this model in the following.

#### 3.2.2. Progressing wound edge guides axonal regrowth after spinal cord injury

In Figs. 8 and 9, we investigate axon growth patterns for a particular ECM stiffness pattern that is intended to represent a stiff, progressing wound edge (see Fig. 4) for two different cone angles of *α* = 30° and *α* = 75°, respectively, using different sets of mechanosensing parameters *β*_link_ = [2.5, 7.5, 12.5] and *n* = [0.5, 2.5, 5.0]. For *n* = 0.5, both figures exhibit comparatively straight axonal growth trajectories, with a decreasing amount of axonal progression towards the top and bottom edges of the domain with increasing *β*_link_ (top rows in Fig. 8 and Fig. 9). With increasing *n*, axonal trajectories become progressively more tortuous (middle and bottom rows). In addition, with increasing *β*_link_ and *n*, the constriction of axons within the lesion core intensifies. Comparing simulations with cone angles of 30° and 75° at *n* = 0.5 reveals a higher degree of axon bundling for the larger cone angle. Consistently, the number of axons crossing the dorso–ventral or ventro–dorsal axis increases for larger cone angles and higher mechanosensing range parameters *n* (see Tab. 1).

**Figure 8:**
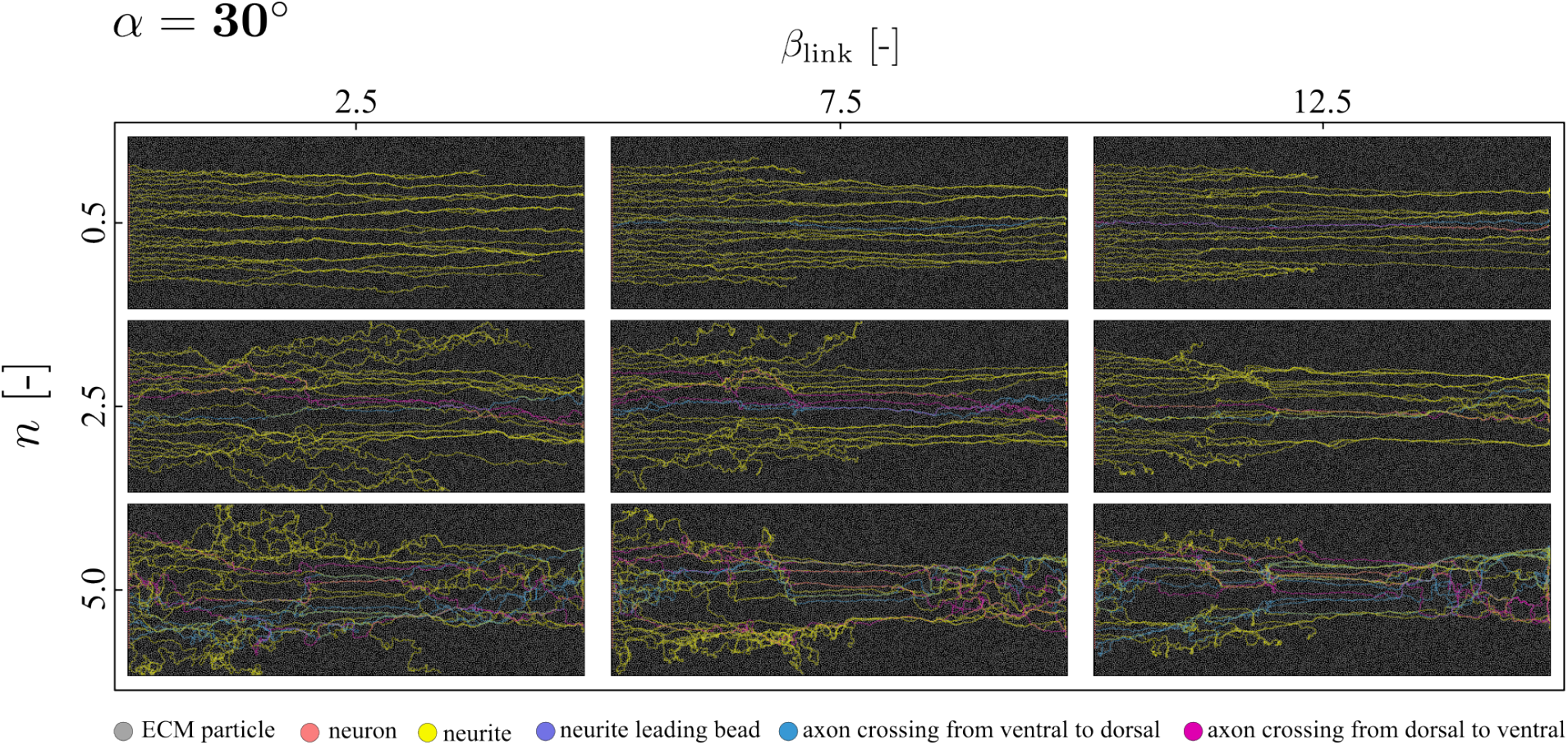
Exemplary agent-based simulation results of axonal regrowth through a stiffness gradient generated by a stiff, progressively evolving wound edge for a cone angle of *α* = 30°. Simulations are shown for mechanosensing parameters *β*_link_ = [2.5, 7.5, 12.5] (increasing from left to right) and *n* = [0.5, 2.5, 5.0] (increasing from top to bottom). For *n* = 0.5, axonal growth remains comparatively straight across all values of *β*_link_, while an increasing *β*_link_ leads to a reduced number of axons progressing near the dorsal and ventral boundaries (top row). The legend in the bottom depicts particle types.

**Figure 9:**
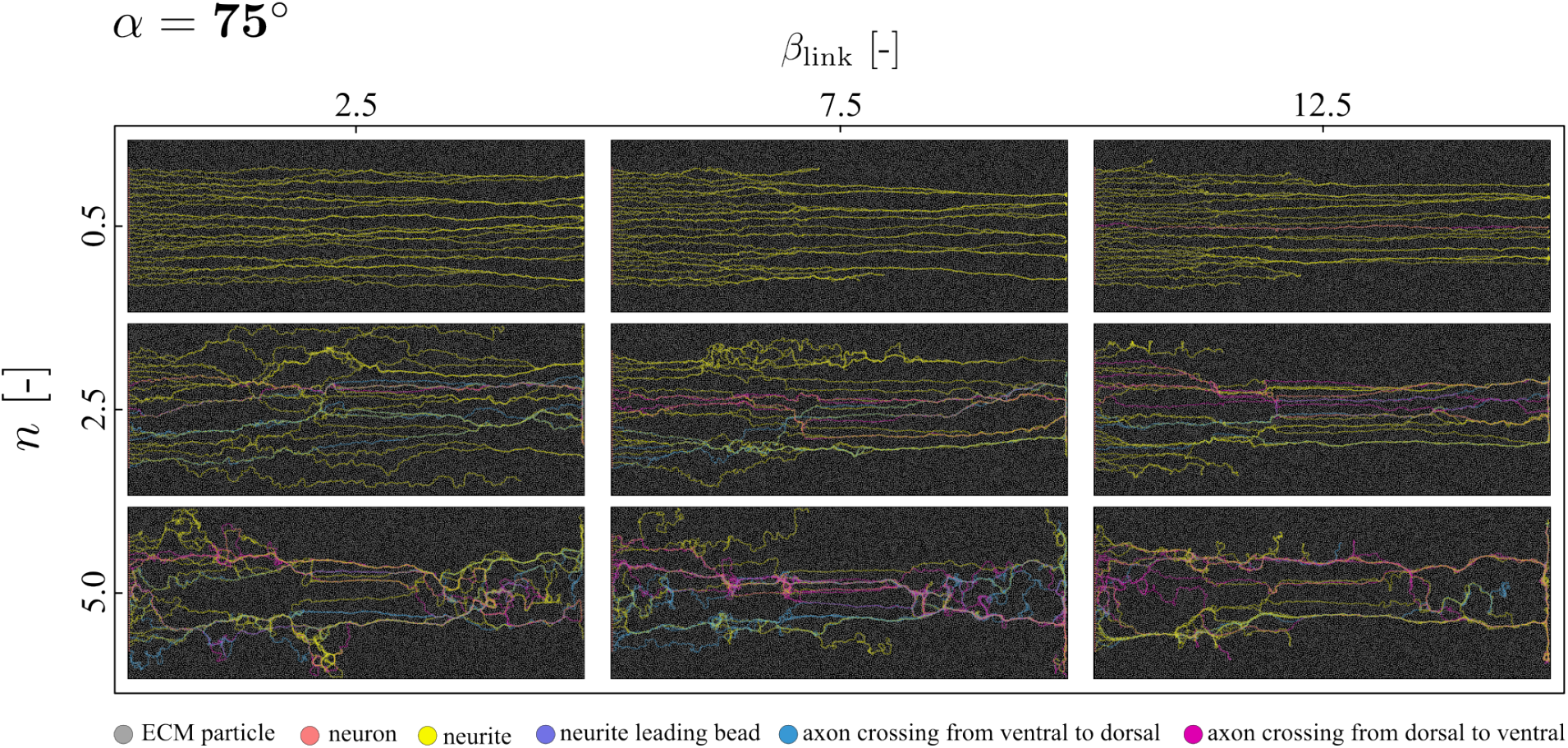
Exemplary agent-based simulation results of axonal regrowth through a stiffness gradient generated by a stiff, progressively evolving wound edge for a cone angle of *α* = 75°. Simulations are shown for mechanosensing parameters *β*_link_ = [2.5, 7.5, 12.5] (increasing from left to right) and *n* = [0.5, 2.5, 5.0] (increasing from top to bottom). For *n* = 0.5, axonal growth remains comparatively straight across all values of *β*_link_, accompanied by a decreasing amount of axonal progression near the dorsal and ventral boundaries with increasing *β*_link_ (top row). The legend in the bottom depicts particle types.

**Table 1:** Summary of rerouting axons (mean ± std) per configuration.

| $\alpha$ | $n$ | $\beta_{\text{link}}$ | $\mathbf{V} \rightarrow \mathbf{D}$ | $\mathbf{D} \rightarrow \mathbf{V}$ |
| --- | --- | --- | --- | --- |
| $30^\circ$ | 0.5 | 2.5 | $1 \pm 1$ | $1 \pm 1$ |
| | | 7.5 | $1 \pm 1$ | $0 \pm 0$ |
| | | 12.5 | $1 \pm 1$ | $0 \pm 1$ |
| | 2.5 | 2.5 | $2 \pm 2$ | $3 \pm 2$ |
| | | 7.5 | $1 \pm 1$ | $2 \pm 1$ |
| | | 12.5 | $1 \pm 1$ | $1 \pm 1$ |
| | 5.0 | 2.5 | $4 \pm 2$ | $3 \pm 1$ |
| | | 7.5 | $5 \pm 1$ | $3 \pm 2$ |
| | | 12.5 | $3 \pm 3$ | $5 \pm 2$ |
| $75^\circ$ | 0.5 | 2.5 | $0 \pm 0$ | $0 \pm 1$ |
| | | 7.5 | $0 \pm 1$ | $1 \pm 1$ |
| | | 12.5 | $0 \pm 0$ | $1 \pm 1$ |
| | 2.5 | 2.5 | $3 \pm 2$ | $4 \pm 2$ |
| | | 7.5 | $5 \pm 2$ | $4 \pm 4$ |
| | | 12.5 | $2 \pm 3$ | $5 \pm 1$ |
| | 5.0 | 2.5 | $5 \pm 2$ | $4 \pm 2$ |
| | | 7.5 | $4 \pm 3$ | $5 \pm 3$ |
| | | 12.5 | $6 \pm 4$ | $5 \pm 3$ |

Tab. 1 summarizes the statistics of rerouting axons crossing the dorso–ventral or ventro–dorsal axis, averaged over five simulations for each combination of the mechanosensing parameters *β*_link_ and *n* at cone angles of 30° and 75°. Overall, simulations with *α* = 30° exhibit slightly fewer rerouting axons than those with *α* = 75°. In addition, an increasing number of rerouting axons is observed with increasing mechanosensing range parameter *n*.

Fig. 10 and Fig. 11 show the tortuosity and mean squared displacement (MSD) as functions of time, averaged over five simulations for all combinations of the mechanosensing parameters *β*_link_ and *n*, and the cone angle *α*. The MSD curves in Fig. 10 exhibit comparable temporal evolutions for both cone angles, *α* = 30° and 75°. In addition, simulation variability increases with increasing values of *β*_link_ and *n*, represented by higher standard deviations. A similar trend is reflected in the tortuosity evolution shown in Fig. 11. The curves for *n* = 0.5 and *n* = 2.5 for both cone angles show rather small degrees of standard deviations across simulations, whereas the results for *n* = 5.0 (last row of Fig. 11) shows both higher absolute values for tortuosity as well as a higher degree of variability across simulations for each individual parameter set. In addition, we can observe rather straight growth directly after each time step of wound-edge evolution, at 25 and 40 % of the total computation time of each simulation, with a decreasing slope in the course of tortuosity over time. This indicates that once the wound edge is developing horizontally, new stiff growth pathways emerge, which result in an increase in the number of straight growing axons, thus reducing the rate at which tortuosity increases over time.

**Figure 10:**
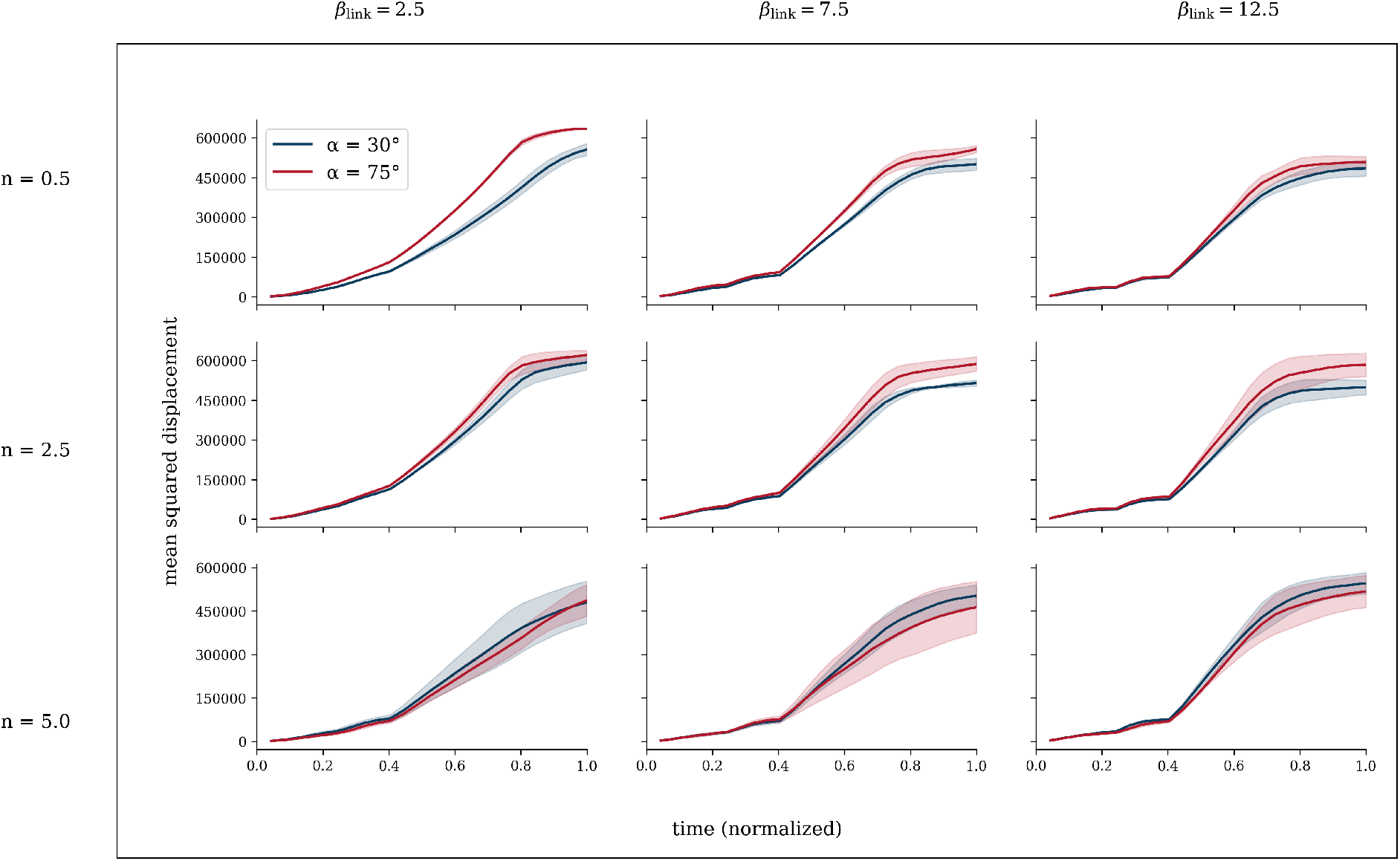
Mean squared displacement (MSD) as a function of time, averaged over five simulations, for all combinations of the mechanosensing parameters *β*_link_ and *n* and for cone angles *α* = 30° (blue) and *α* = 75° (red). Parameter values increase from left to right for *β*_link_ and from top to bottom for *n*. The MSD curves exhibit comparable temporal evolution for both cone angles, while increasing values of *β*_link_ and *n* lead to higher variability between simulations, reflected by increasing standard deviations (shown in shaded regions). Time was normalized such that 1 corresponds to 100% of the ABM simulation’s computation time.

**Figure 11:**
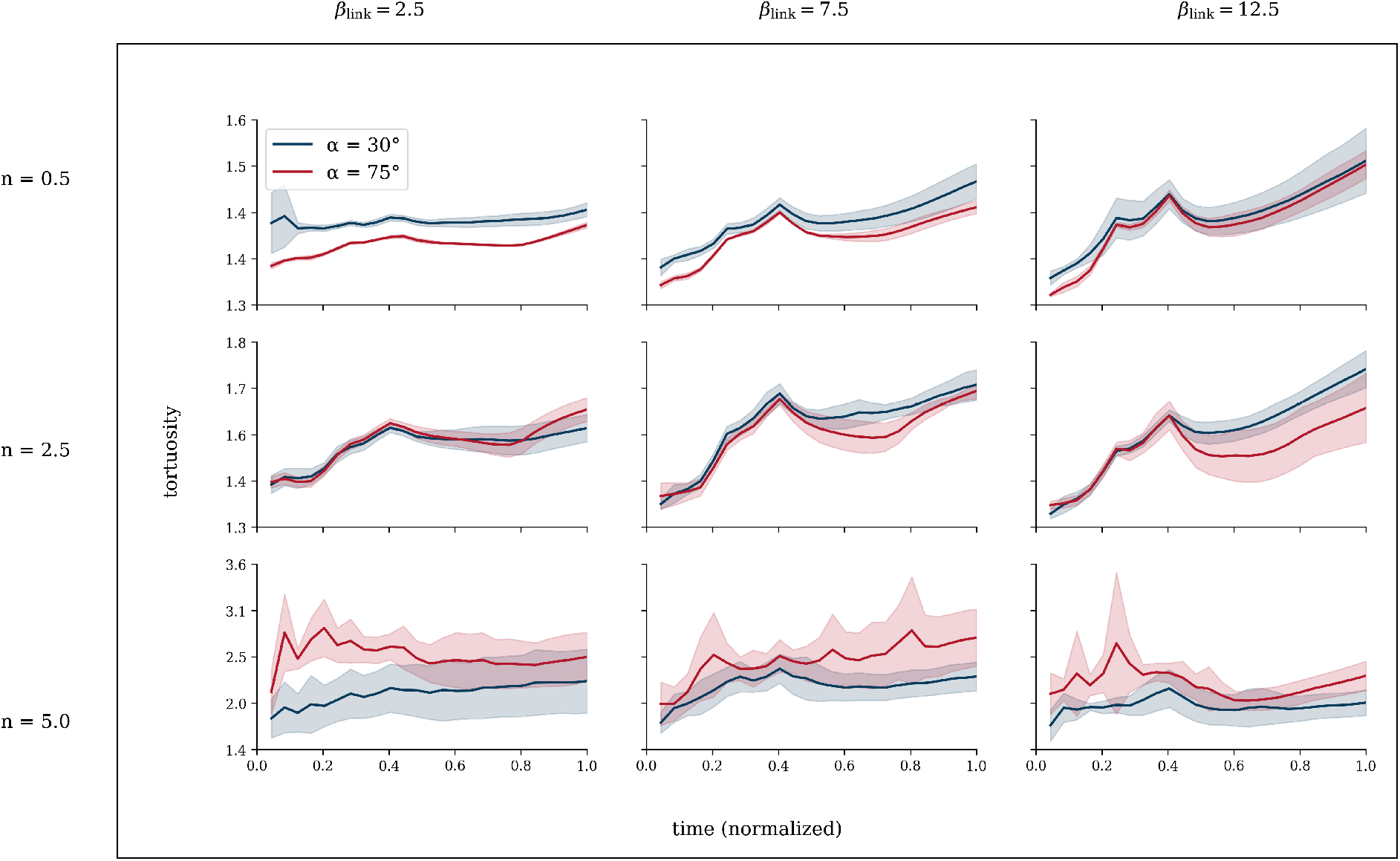
Tortuosity as a function of time, averaged over five simulations, for all combinations of the mechanosensing parameters *β*_link_ and *n* and for cone angles *α* = 30° (blue) and *α* = 75° (red). Parameter values increase from left to right for *β*_link_ and from top to bottom for *n*. For *n* = 0.5 and *n* = 2.5, tortuosity exhibits relatively low variability across simulations for both cone angles, whereas *n* = 5.0 (bottom row) results in higher absolute tortuosity values and increased variability. Straight growth is observed immediately following each wound-edge evolution step (at 25% and 40% of the total simulation time), indicated by reduced slopes in tortuosity over time, suggesting that horizontally evolving wound edges promote the emergence of straighter growth pathways. Standard deviations are shown in shaded regions. Time was normalized such that 1 corresponds to 100% of the ABM simulation’s computation time.

To further assess the capabilities of the proposed model, we compared individual microscopic images of zebrafish larvae recorded throughout regeneration (Figs. 5 and 6) with simulation outputs at corresponding stages of the computational time course. Figure 12 summarizes representative cases, in which the collective organization of individual axonal trajectories qualitatively agrees between experiment and simulation. The examples are arranged by regeneration stage: the top row shows early regeneration alongside simulations at 25 % of the total computation time, the middle row depicts intermediate regeneration matched to 40 % of the computation time, and the bottom row highlights successfully regenerated larvae together with results at 100 % (end) of the simulation. To enhance visual comparability, microscopic images of zebrafish that have not yet fully regenerated are cropped in the top and middle rows to match the corresponding simulation views.

**Figure 12:**
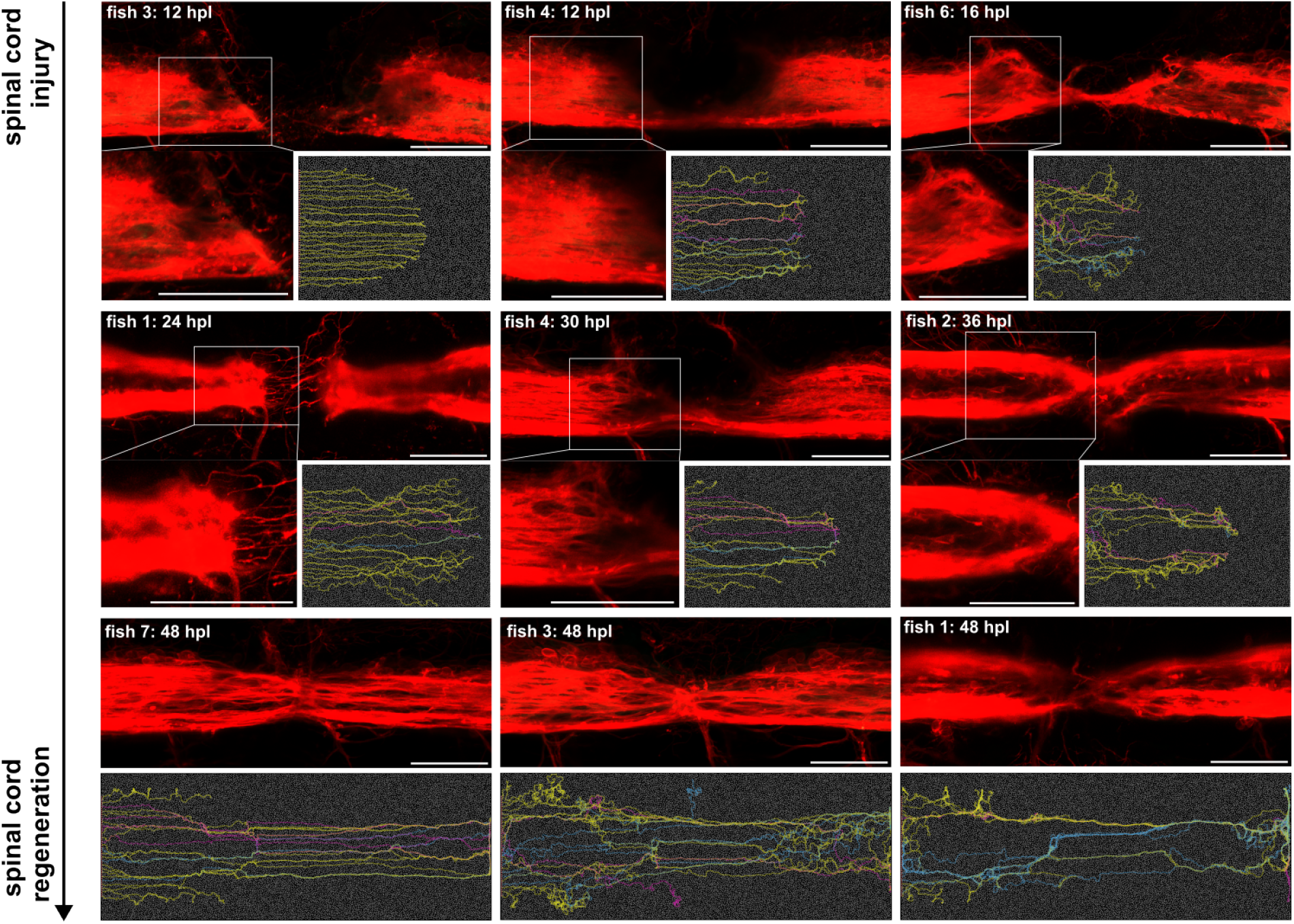
Exemplary experimental images of zebrafish at different time stamps during regeneration with qualitatively matching agent-based model simulation results and associated parameters (*β*_link_, *n, α*) and simulation progress percentage. Fish 3 (12 hpl) (12.5, 0.5, 30°) at 25%; Fish 4 (12 hpl) (2.5, 2.5, 75°) at 25%; Fish 6 (16 hpl) (2.5, 5.0, 30°) at 25%; Fish 1 (24 hpl) (2.5, 2.5, 30°) at 40%; Fish 4 (30 hpl) (12.5, 2.5, 75°) at 40%; Fish 2 (36 hpl) (12.5, 5.0, 30°) at 40%; Fish 7 (48 hpl) (12.5, 2.5, 75°) at 100%; Fish 3 (48 hpl) (7.5, 5.0, 30°) at 100%; Fish 1 (48 hpl) (12.5, 5.0, 75°) at 100%. Scale bars: 50 *µ*m.

We observe that multiple parameter combinations (*β*_link_, *n, α*) can reproduce the overall growth patterns seen *in vivo*. Overall, simulations with the smallest mechanosensing exponent *n* = 0.5, which controls how strongly local stiffness modulates the growth directionality, show weaker qualitative agreement with the experimental images. Beyond this, no consistent trend with respect to *n* is apparent. In contrast, the growth speed parameter *β*_link_ exhibits a clearer stage dependence: lower values (*β*_link_ = 2.5) tend to better capture early regeneration patterns (Fish 4, 12 hpl; Fish 6, 16 hpl; Fish 1, 24 hpl), whereas higher values (*β*_link_ = 12.5) more closely resemble later stages beyond the first day (Fish 4, 30 hpl; Fish 2, 36 hpl; Fish 7, 48 hpl; Fish 1, 48 hpl). Finally, both tested growth-cone angles, *α* ∈ {30°, 75°}, are able to reproduce the observed microscopic axon trajectory patterns.

## 4. Discussion

The computational model presented in this study offers an approach to qualitatively compare agent-based axonal growth simulations with time-lapse confocal microscopy images of axonal regrowth during spinal cord regeneration in zebrafish larvae. Based on the assumption of a stiffer progressing wound edge, mechanosensing-driven growth parameters were employed to guide axonal regrowth such that globally emerging patterns are reconstructed. This model can be used to test assumptions and hypotheses based on experimental investigations like [7, 3, 6, 17] which implicate mechanics in neurodevelopment and axon regeneration. As this field of research is still at an early stage, many of the underlying coupled biochemical and biophysical mechanisms require further investigation to enhance interpretation of the interplay between mechanics, biology and biochemistry during axon (re)growth. Experimental investigations that probe effects and correlations in these different disciplines are increasingly complex and difficult to perform, particularly *in vivo* [15]. Here, we propose to combine experimental data with *in silico* modeling to explore to which extent different mechanisms of mechanosensing can reconstruct axonal growth patterns observed *in vivo* (Fig. 7).

Our findings show that a suitable set of growth-governing parameters together with biologically inspired stiffness gradients can qualitatively reproduce global patterns that naturally emerge over the course of regeneration in zebrafish larvae (see Fig. 5 and 6). Based on repeatedly observed growth patterns, we conjecture a progressively evolving wound edge which forms a stiffer mechano-attractant for axonal pathfinding. A similar mechanism was experimentally observed by Shellard et al., who showed that neural crest cells in embryonic development self-generate a stiffness gradient in the adjacent tissue (similar to our approach) which acts as a durotactic guidance cue for cell migration [20]. Although species and neurobiological process in this study differ from those in our computational approach (spinal cord regeneration in zebrafish larvae), we propose that the *in vivo* paradigm of an evolving edge that creates a stiffness gradient in the surrounding tissue of migrating neuronal cells [20] can also be leveraged to reproduce the global patterns of regenerating axons *in silico* (Fig. 12). Based on these findings, new experimental investigations can test whether stiffness gradients assumed in this study, a transient stiff wound edge progressively evolving through the lesion core, are actually present in the spinal cord of the regenerating zebrafish larvae. This will enable an iterative feedback loop where computational modeling can be refined based on experimental findings and experiments can be redesigned based on computational outcomes.

We observe that the simulations reproduce the experimentally observed axonal growth trajectories for both investigated cone angles (*α* = [30, 75]°). The mechanosensing parameter *n*, which regulates the coupling of growth directionality to environmental stiffness, strongly influences the resulting final growth patterns, with *n* = 5.0 yielding the closest qualitative agreement with experimental observations. In contrast, the mechanosensing parameter *β*_link_, governing the coupling of growth speed to environmental stiffness, exhibits a comparatively lower influence. This suggests that, within the current modeling framework (employing a power-law formulation for *n* and a logistic evolution for *β*_link_), stiffness-modulated growth speed plays a more dominant role than directionality in simulating the overall axonal pathfinding outcome after spinal cord regeneration in zebrafish.

In the context of the influence of environmental stiffness on neuronal growth speed, experimental results reported by Kravikass et al. indicate no significant change in the instant growth velocity of rodent hippocampal neurons across hydrogels of varying stiffness in 3D [19]. The effective velocity (end-to-end distance divided by trajectory duration) does increase with stiffness in this study. However, this reflects the higher directional persistence of growth in stiffer matrices rather than an increase in actual growth speed. Although this confirms our second assumption illustrated in Fig. 2c, it is inconsistent with the hypothesis of accelerated growth in a stiffer environment. Similarly, Koser et al. report that growth cone motility of *Xenopus laevis* retinal ganglion cells was, if anything, slightly higher on softer substrates *in vitro* [7]. The higher axon extension velocity reported on stiffer substrates in the same study likewise reflects straighter, more persistent growth rather than faster traversal of the growth cone along its path. Interestingly, the same study also observed that axon bundles turn toward softer regions when growing on stiffness gradients *in vivo*. As proposed in the agent-based model presented in this study, this may arise because axons on the stiffer side of the bundle grow faster, collectively steering the bundle toward softer tissue. In this context, Oliveri et al. developed a mathematical framework proposing that individual growth cones within a growing axon bundle perceive different signals depending on their position within the fascicle, thereby exerting slightly different traction forces on the surrounding substrate [25]. Their mechanical coupling within the bundle gives rise to a net force and torque at the collective level, resulting in changes in bundle growth and turning. In their framework, the ability of growth cones to grip the surrounding medium and generate traction is enhanced on stiffer substrates, while the resultant force affects growth speed and the torque drives bundle reorientation [25]. However, as the study is primarily theoretical, it does not provide experimental evidence to determine whether stronger traction on stiffer substrates necessarily leads to faster individual axon advancement or whether this effect is counterbalanced by a proportional increase in substrate resistance.

When considering different stages of regeneration, however, *β*_link_ emerges as a more important modulator of temporal growth dynamics: lower values of *β*_link_ better replicate early regeneration stages (12–24 hours post lesion), whereas higher values more closely resemble axonal growth patterns observed at later stages (30–48 hours post lesion).

The presented approach combines simulations of individual axons governed by mechanically inspired growth paradigms with the analysis of globally emerging growth patterns arising from interactions between multiple growing axons, highlighting the versatility of the modeling framework. This multiscale perspective enables direct comparison with specific experimental observations reported in the literature. For instance, it has been shown that regenerating descending axons in the adult zebrafish reroute from white to grey matter tissue [26], and that a subset of descending axons crosses from the ventral to the dorsal side during spinal cord regeneration of the zebrafish larvae [27]. Quantitatively, [27] reports that approximately 40 % of regenerating descending axons switch from ventral to dorsal trajectories which is comparable to our simulation results yielding a maximum of approximately 42 % ventro–dorsally rerouting axons for the parameter set (*β*_link_, *n, α*) = (12.5, 2.5, 75°) (bottom row in Tab. 1: 10 out of 26 regrowing axons, representing the upper end of the observed range). Other parameter combinations, however, produce lower fractions of ventro–dorsal crossings. In contrast to the experimental observations reported in [27], indicating little to no regeneration on the dorsal side, the present simulations tend to yield more symmetric growth patterns, with comparable numbers of dorso–ventral and ventro–dorsal crossings, pointing to potential limitations of the current modeling assumptions. In addition, another growth mechanism reported in the literature emerges naturally from the simulations, namely the bundling of individual axons into fascicles, a phenomenon that has been repeatedly observed *in vivo* [24, 28, 29]. After not observing durotaxis on physiological stiffness gradients for single cells, [20] suggested that durotaxis could be interpreted as emerging property from this fasciculation. This interpretation is consistent with our simulation results, which show that a more accurate representation of the ECM stiffness landscape leads to more pronounced collective outgrowth into stiffer regions. Furthermore, with increasing parameters *β*_link_ and *n*, the resulting growth patterns exhibit improved qualitative agreement with the microscopic observations reported in this study. Moreover, higher degrees of these simulated axonal fasciculation, which correspond to larger mechanosensing parameter values, are associated with an increased fraction of ventro–dorsally crossing axons in the simulations which aligns more closely with the experimental findings reported in [27].

### 4.1. Limitations

A limitation of this study may be that the model is based on a relatively small experimental dataset obtained from zebrafish larvae undergoing repeated embedding and imaging procedures.

As a consequence, experimental images of dorsal and lateral views were acquired at different time points during regeneration, limiting direct temporal comparability between these perspectives.

For the sake of model simplicity, the parameter sensitivity analysis focused exclusively on mechanosensing-related parameters, while other parameters such as the steepness of the stiffness gradients, particle density or similar where not systematically explored.

The mechanisms underlying the progressive ventro–dorsally filling observed in the lateral view during regeneration are not yet fully understood and are therefore not explicitly captured in the current simulations.

The presented images are maximum-intensity projections of z-stacks acquired across different depths (medio–lateral for Fig. 5 and dorso–ventral for Fig. 6). Consequently, the apparent two-dimensional trajectories reflect projected three-dimensional paths, and the exact courses of individual axons cannot be reconstructed from these data alone.

The mathematical functions chosen to introduce mechanosensing into the model, a logistic function coupling growth rate to stiffness and a power law coupling link range to stiffness, have been shown to successfully recreate the experimentally observed global growth patterns. However, other, more biophysically motivated descriptions may prove even more suitable and physically interpretable. Finally, we acknowledge that the currently available experimental evidence on the relationship between axon growth dynamics and environmental stiffness [7, 19] does not support one of the two assumptions used in our simulations, namely faster growth on stiffer substrates. Rather, these studies more strongly suggest that increased stiffness promotes greater directional persistence, whereas the pathwise growth speed itself does not appear to increase accordingly. We suspect that extending the current 2D model to 3D, incorporating a more realistic grey- and white-matter morphology of the zebrafish spinal cord together with corresponding stiffness profiles, and including additional guidance mechanisms such as chemotaxis could resolve this discrepancy, potentially rendering an explicit coupling between stiffness and growth speed unnecessary while still reproducing the experimentally observed growth patterns. Nevertheless, to the best of our knowledge, there is currently no direct *in vivo* experimental evidence from the zebrafish spinal cord that definitively rules out this assumption. As the primary aim of this study is to establish a computational framework for investigating possible mechanisms of mechanosensing during axonal regrowth after spinal cord injury in zebrafish, we defer this question to future work where we plan to extend the model and test it against dedicated *in vivo* regeneration data.

## 5. Concluding Remarks

Further development of the model presented here will enable the investigation of additional growth paradigms and potential mechanisms by which mechanical and chemical cues guide axonal regeneration. In particular, we will extend the framework to three dimensions and incorporate a two-material representation of the spinal cord, together with more realistic axonal growth initiations. This would enable a more detailed computational analysis to test specific hypotheses. For instance, Becker et al. observed that axons reroute from the rostral white matter into the caudal grey matter during regeneration of the adult zebrafish spinal cord [26]. Moreover, further investigation of optimal stiffness ranges for axonal regrowth is warranted, as experimental evidence suggests the existence of a specific window of substrate stiffness that promotes regeneration, while both lower and higher stiffness values are less permissive. Koch et al. demonstrated that in the peripheral nervous system dorsal root ganglion neurons exhibit the largest axonal extensions on substrates with a stiffness of approximately 1000 Pa, compared with substrates spanning stiffness values between 150 and 5000 Pa [30]. In addition, Kayal et al. demonstrated *in vitro* that neuronal elongation and orientation of NG108-15 neural cells cultured on hydrogels are influenced by both the absolute substrate stiffness and the steepness of the imposed stiffness gradient [16]. Building on these observations, it would be highly informative to further investigate how these two parameters jointly affect the globally emerging axonal growth patterns in the present modeling framework. Addressing such questions in a 3D *in silico* larval model could shed further light on the role of tissue mechanics in guiding spinal cord regeneration. Furthermore, Pillai et al. reported a functional interplay between mechanical and chemical signaling during brain development in *Xenopus laevis* [31]. Incorporating additional biochemical fields into the present modeling framework, and coupling these fields to the mechanical interactions between growing axons and their environment, could therefore further extend the predictive and explanatory capabilities of the proposed approach.

The model approach presented here can contribute to a deeper understanding of the mechanisms underlying spinal cord regeneration and, in the longer term, help inform therapeutic strategies by facilitating the translation of these insights to mammalian systems and, ultimately, to humans following spinal cord injury.

## 6. Acknowledgements

The funding from the German Research Foundation (Deutsche Forschungsgemeinschaft, DFG) through project 460333672 CRC1540 EBM (project B01) is gratefully acknowledged. The authors gratefully acknowledge the agent-based modeling framework developed within project C01, as well as the confocal live imaging data of zebrafish spinal cord regeneration provided by project B05. This work was supported by the Imaging Facility and the Animal Facility of the Max-Planck-Zentrum für Physik und Medizin.

## Declaration of AI and AI-assisted technologies in the writing process

During the preparation of this work the author(s) used Deepl and ChatGPT in order to improve readability and language. After using this tool/service, the author(s) reviewed and edited the content as needed and take(s) full responsibility for the content of the publication.

