## Supplementary Material for "*In silico* model of axonal pathfinding during spinal cord regeneration in zebrafish larvae"

Table S1: Computational parameters used for the test-case simulations shown in Fig. 7. All simulations were performed for both cone angles $\alpha = 30°$ and $\alpha = 75°$, while durotaxis parameters were varied as indicated.

| **Category** | **Parameter** | **Value** |
|---|---|---|
| Simulation | Pixel size $p_x$ | 0.5 |
| | Time step $\Delta t$ | 0.002 |
| | Total simulation time | 30 |
| Neuron properties | Neuron radius | 1.25 |
| | Neuron mobility | 0.7 |
| | Neurite radius | 0.5 |
| | Neurite mobility | 1.0 |
| | Neuron layer index | 0 |
| Growth mechanics | Straight growth | 1 |
| | Growth rate | 10 |
| | Maximum neurite length $l_{\text{max}}$ | 1.2 |
| | Cone angle $\alpha$ | 0.523599 |
| | Link range | 30 |
| Domain geometry | Domain size $(L_x, L_y)$ | (200, 200) |
| Spring constants | $k_{\text{low}}$ | 0.01 |
| | $k_{\text{med}}$ | 1.0 |
| | $k_{\text{high}}$ | 4.0 |
| | $\varepsilon$ | 2.0 |
| ECM properties | ECM volume fraction | 0.5 |
| | ECM particle radius | 1.0 |
| | ECM mobility | 0.5 |
| | Link reset time | 0.01 |
| Durotaxis | Durotaxis cone angle | 0 |
| | Durotaxis link range | 0 |
| | Durotaxis parameter | 1 |
| | Durotaxis sensing range | 30 |
| Stiffness patterning | Stiffness factor | 2.5 |

Table S2: Computational parameters used for simulations with a progressively evolving wound-edge stiffness gradient. All simulations were performed for both cone angles $\alpha = 30°$ and $\alpha = 75°$, while durotaxis parameters $\beta_{\text{link}}$ and $n$ were systematically varied.

| **Category** | **Parameter** | **Value** |
|---|---|---|
| Simulation | Pixel size $p_x$ | 0.5 |
| | Time step $\Delta t$ | 0.002 |
| | Total simulation time | 300 |
| Neuron properties | Neuron radius | 1.25 |
| | Neuron mobility | 0.7 |
| | Neurite radius | 0.5 |
| | Neurite mobility | 1.0 |
| | Neuron layer index | 0 |
| Growth mechanics | Straight growth | 0 |
| | Growth rate | 10 |
| | Maximum neurite length $l_{\text{max}}$ | 1.2 |
| | Cone angle $\alpha$ | 0.523599 |
| | Link range | 30 |
| Domain geometry | Domain size $(L_x, L_y)$ | (800, 300) |
| Spring constants | $k_{\text{min}}$ | 0.01 |
| | $k_{\text{med}}$ | 1.0 |
| | $k_{\text{max}}$ | 4.0 |
| | $\varepsilon$ | 2.0 |
| ECM properties | ECM volume fraction | 0.5 |
| | ECM particle radius | 1.0 |
| | ECM mobility | 0.5 |
| | Link reset time | 0.01 |
| Durotaxis | Durotaxis cone angle flag | 1 |
| | Durotaxis link range flag | 1 |
| | Durotaxis activation parameter | 1 |
| | Durotaxis sensing range | 30 |
| | $\beta_{\text{link}}$ | 2.5 |
| | $n$ | 0.5 |
| Stiffness patterning | Stiffness factor | 2.5 |

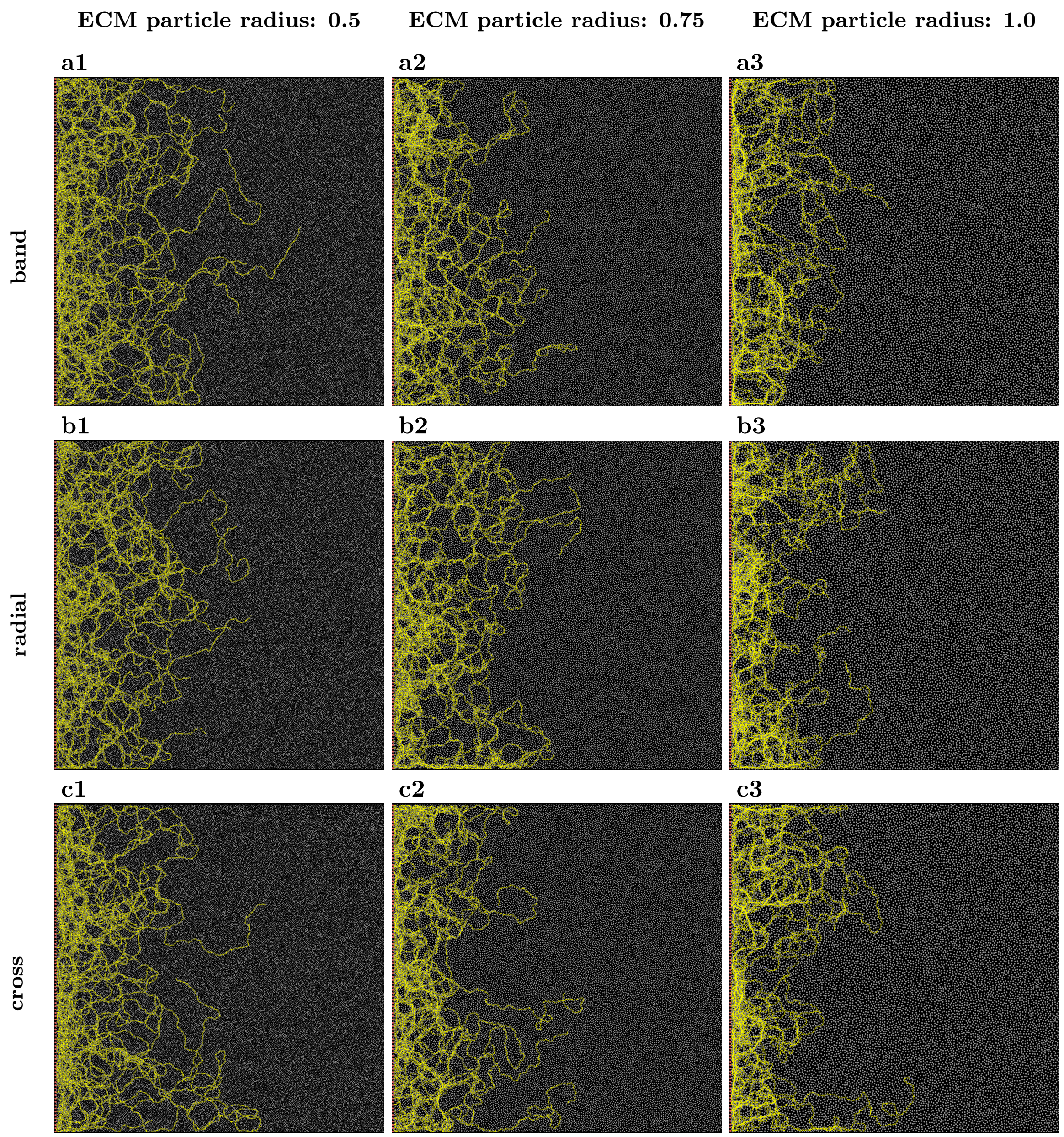


Fig. S1: Effect of the size of ECM particles on axonal growth patterns for three representative test cases. Columns correspond to increasing radii of 0.5, 0.75, and 1.0. Rows show (a) banded, (b) radial, and (c) cross-shaped stiffness configuration.
